# Growth-induced collective bending and kinetic trapping of cytoskeletal filaments

**DOI:** 10.1101/2024.01.09.574885

**Authors:** Deb Sankar Banerjee, Simon L. Freedman, Michael P. Murrell, Shiladitya Banerjee

## Abstract

Growth and turnover of actin filaments play a crucial role in the construction and maintenance of actin networks within cells. Actin filament growth occurs within limited space and finite subunit resources in the actin cortex. To understand how filament growth shapes the emergent architecture of actin networks, we developed a minimal agent-based model coupling filament mechanics and growth in a limiting subunit pool. We find that rapid filament growth induces kinetic trapping of highly bent actin filaments. Such collective bending patterns are long-lived, organized around nematic defects, and arises from competition between filament polymerization and bending elasticity. The stability of nematic defects and the extent of kinetic trapping are amplified by an increase in the abundance of the actin pool and by crosslinking the network. These findings suggest that kinetic trapping is a robust consequence of growth in crowded environments, providing a route to program shape memory in actin networks.

## Introduction

Actin networks organize into diverse architectures and morphologies within cells, in order to regulate vital physiological functions such as cell motility, cell division, and endocytosis (1–4). Observed actin-based structures include linear actin arrays in stress fibers and filopodia (5–8), branched networks in lamellipodia (4, 9), actomyosin rings (10), and disordered networks (11). Achieving the desired structure poses a challenge for the cell due to the constant reorganization of actin filaments driven by polymerization, disassembly, myosin-based contractility, and crosslinking (2, 4, 12). Moreover, intracellular crowding impacts the assembly of desired architectures. While previous research has predominantly focused on self-organization of actin networks driven by myosin motors and crosslinking (2, 13–21), the influence of crowding on the growth of actin network structures remains poorly elucidated.

In this study, we theoretically investigate the growth of actin networks in a limiting subunit pool. Our study is inspired by recent *in vitro* experiments of F-actin growth from a limiting pool of monomers and nucleators (22). The limitation of monomer abundance constrains the assembly of individual filaments by pool depletion and can be utilized to achieve filament length control (23, 24). Additionally, the abundance of nucleators provides a means to regulate both filament length and density. It has been experimentally shown that in sufficiently crowded environments, long actin filaments assemble into highly curved morphologies that act as templates for subsequent assembly of neighboring actin filaments (22). These high curvature filaments are unable to relax stresses by turnover, suggesting a means to encode shape memory in actin filament assemblies.

To understand the physical origin of these observations, we employed dynamic Monte Carlo and Langevin dynamics simulations (18) to model the growth of semi-flexible actin filaments from a limiting monomer pool. In our model, the number of filaments is fixed by the number of nucleators, and filaments interact via volume exclusion and crosslinking. While volume exclusion effects have often been overlooked in active filament simulations, actin networks *in vitro* (22, 25) and within the cortex (26) are constrained within a thin two-dimensional layer, where steric interactions may play a significant role in shaping actin networks (27). By incorporating these effects, our model allows us to explore how the interplay between filament growth, elasticity, and crowding influences the organization of actin networks under limiting subunit resources.

We begin by formulating an effective model for the dynamics of filament length within a finite pool of monomers. Subsequently, we integrate this effective model of filament length control into agent-based simulations to explore the self-organization of growing networks. Our study illustrates how factors such as filament nucleation, actin density, and crosslinker density influence the resulting network morphology. To characterize the emergent network architecture and mechanics, we measure compressive strains, bending energies in filaments, as well as nematic order and defect organization. We observe that an increased number of nucleators promotes kinetic trapping of highly curved filamentous structures, inducing shape memory in actin filament assemblies. The extent of kinetic trapping can be enhanced by increasing actin density and crosslinker concentration. Our findings elucidate how individual filament morphologies can be tuned by altering the network composition, consequently shaping the collective network morphology.

## Results

### Filament growth control in a limiting monomer pool

Actin filaments grow via assembly and disassembly of G-actin monomers at both ends. Within the cellular environment, various actin-binding proteins govern different aspects of actin filament growth. Recent studies show that the cellular actin structures compete for actin monomers, and that their range and size are governed by the size of the monomer pool (24). This limiting pool scenario is also applicable to *in vitro* reconstitution assays (22). It has been shown previously that the steady-state length distribution of actin filaments grown from a restricted monomer pool is exponential, lacking a characteristic filament length (28, 29). However, the dynamics of filament length over physiologically relevant timescales (ranging from minutes to hours) depends on several other factors and remains less elucidated.

To investigate the collective dynamics of growing filaments, we first formulated an effective model for the growth of a single filament within a limiting monomer pool. We model individual actin filament growth from a shared limiting pool of actin monomers using a Dynamic Monte Carlo scheme (see Methods for details) (Fig. 1A). Assuming no spontaneous nucleation occurs, the number of growing filaments is set by the number of nucleators chosen at the beginning of the simulation. We find that over the timescales of a few hours, the fluctuation in actin filament length depends on the size of the monomer pool. For small pool size, the filaments exhibit large length fluctuations (Fig. 1B), and lack a typical length as the size distribution is exponential (Fig. 1C). However, with a sufficiently large monomer pool, these fluctuations occur over much longer timescales (beyond hours), resulting in filaments growing to similar lengths, with a well-defined mean length (Fig. 1B & C). In this large monomer pool size limit, the dynamics of filament length is given by

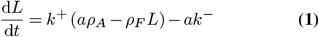

where *L*(*t*) is the filament length at time *t, a* is the monomer size, *k*^±^ are the assembly and disassembly rate constants, *ρ*_*A*_ = *N/V* is the total actin density, and *ρ*_*F*_ = *N*_*F*_ */V* is filament density. The number of filaments, total number of monomers (pool size) and system size are given by *N*_*F*_, *N* and *V*, respectively. In this effective model, the timescale to reach steady-state is given by *T*_*s*_ = (*k*^+^*ρ*_*F*_)^−1^.

**Fig. 1.**
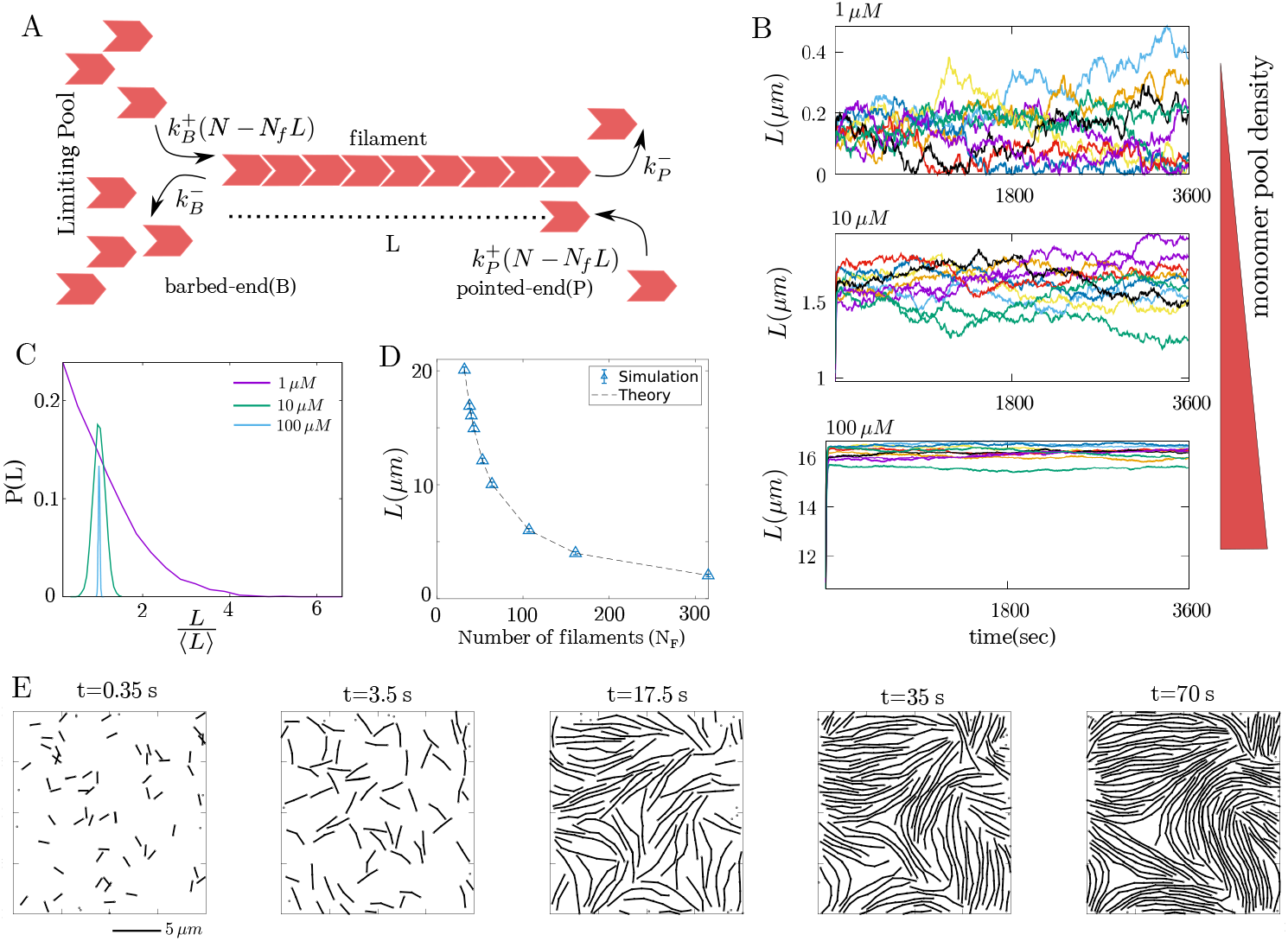
Filament length control in a limiting monomer pool. (A) Schematic showing the model for actin filament growth within a limiting pool of monomers (B) Fluctuation in filament length over one hour for three different monomer pool sizes: 1 *μ*m (top), 10 *μ*m (middle), and 100 *μ*m (bottom). (C) Filament length distribution at different monomer pool sizes. (D) Mean filament length as a function of number of filaments (*N*_*F*_). (E) Configurations of a collection of growing filaments at (left to right) *t* = 0.35s, 3.5s, 17.5s, 35s, and 70s. Parameters values used here are *k*^+^ = 11.4*/μM s, k*^*−*^ = 1*/s* and *N*_*F*_ = 10 for different pool sizes in B and C. For panel D, [actin]=10 *μM* and for panel E, [actin]=15 *μ*M, and *N*_*F*_ = 68.

Setting *dL/dt* = 0 yields the steady-state filament length as 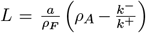, in excellent agreement with the stochastic simulations of the full model (Fig. 1D). Consequently, we use (Eq. 1) to model the length dynamics of actin filaments in agent-based simulations (Fig. 1E), to better understand the impact of filament growth on network formation over the timescale of a few hours.

### Growth-induced bending and kinetic trapping

We incorporated the single-filament growth model described in the preceding section into AFINES, a simulation framework for cytoskeletal networks (18) (see Methods for details). In these simulations, filament growth was initiated from small “seeds” (nucleators) within a quasi-two-dimensional environment with volume exclusion interaction, as shown in Fig. 1E. By maintaining a constant density of total actin monomers, the filament length was determined by the nucleator number density, with a higher nucleator density resulting in shorter filaments (Fig. 1D). The changes in filament length significantly influenced the emergent morphology of the growing network (Fig. 2A), leading to actin networks with varying degrees of nematic order, topological defects, and filament curvatures (see Movies S1-S4).

**Fig. 2.**
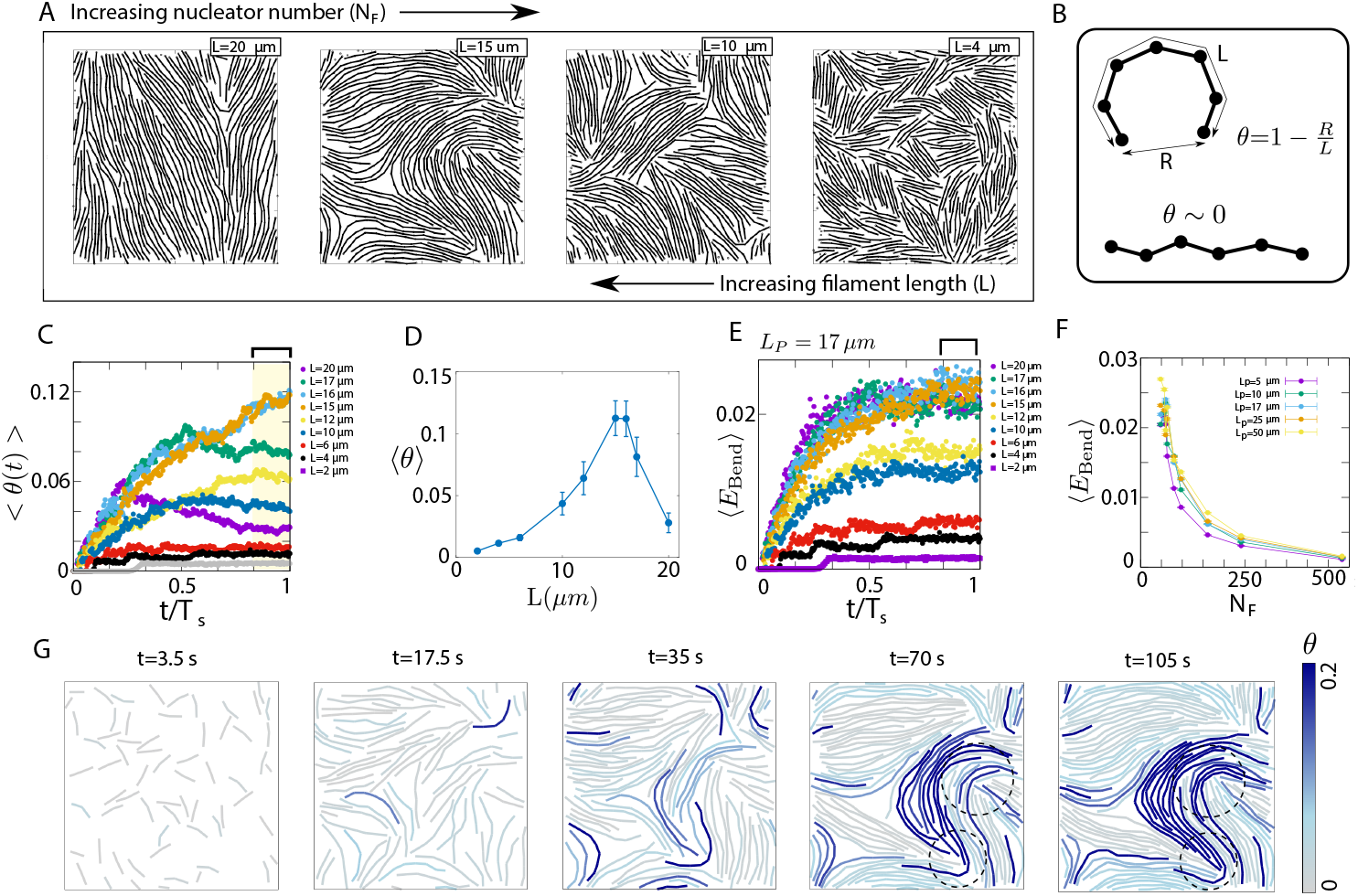
Collective bending pattern and kinetic trapping of filaments. (A) Configurations of actin filament assemblies for different nucleator densities (i.e., different filament lengths). (B) Quantification of filament bending by the compressive strain *θ*. (C) Time evolution of average compressive strain per filament, *θ*(*t*), for different filament lengths. (D) Steady-state value of the ensemble-averaged compressive strain per filament, for different filament lengths. (E) Dynamics of the average bending energy per filament, for different filament lengths. (F) Average bending energy for different nucleator numbers for varying bending rigidity. (G) Time-lapse of filament configurations in growing networks, with filaments color coded by their compressive strains. Over time we observe the emergence of correlated bending due to kinetic trapping of high curvature filaments (dashed circle). The actin density is 15 *μM* for all results shown here. For panel G, *N*_*F*_ = 64 (*L* = 15 *μ*m).

To assess the mechanical properties of the actin filament assembly, we quantified the extent of filament bending by calculating the filament compressive strain *θ*(*t*) = 1 − *R*(*t*)*/L*(*t*) (Fig. 2B), where *R*(*t*) represents the end-to-end length and *L*(*t*) is the filament contour length at time *t*. As filaments grew and began interacting, the ensemble-averaged compressible strain ⟨*θ*(*t*)⟩ increased over time (Fig. 2C). We mostly observed a monotonic increase in ⟨*θ*(*t*)⟩ over time for filaments of short to intermediate lengths (*L < L*_*p*_). For longer filaments, *L* ≥17 *μ*m, *θ*(*t*) peaked and subsequently relaxed to a lower value, as shown in Fig. 2C. These relaxation events indicate instances of structural relaxation for kinetically trapped filaments with high bending energy, as we discuss in more detail later.

Notably, the steady-state value of the compressive strain ⟨*θ*⟩ exhibited a non-monotonic relationship with the steady-state filament length *L*, reaching its peak at an intermediate filament length (Fig. 2D). At low filament densities, filaments grew to lengths much longer than their persistence lengths and encountered less obstacle to growth due to a lower steric hindrance. This resulted in a lower value of⟨ *θ*⟩ and a high nematic order. Conversely, at high filament number density, filament lengths were smaller, resulting in increased resistance to bending and smaller values of ⟨*θ*⟩. Short filament assemblies also possess a locally high nematic order, with the presence of many domain boundaries (see Fig. 2A-right). At an intermediate filament density, filaments grow to longer lengths and experience buckling due to growth-induced pressure from the surrounding filamentous environment. A similar non-monotonic trend in filament curvature versus length has been observed *in vitro* assays involving actin and Formin, where changing Formin density altered the nematic alignment, defect morphologies and filament curvatures in F-actin networks (22).

Much like the compressive strain, the average bending energy stored per filament, ⟨*E*_Bend_(*t*)⟩, increased over time as filament growth induced collective bending patterns (Fig. 2E). The steady-state value of⟨ *E*_Bend_⟩ decreased with an increase in filament number *N*_*F*_, showing minimal dependence on variations in the bending rigidity of the filaments (as characterized by the persistence length *L*_*p*_) (Fig. 2F). This suggests that changes in nucleator density primarily influence filament bending through alterations in filament length. To understand the origin of collective filament bending and the emergence of high-curvature shapes depicted in Fig. 2A, it is informative to analyze the time-lapse of filament configuration in a growing assembly (Fig. 2G). We observe that filament bending is induced due to growth within a locally crowded filamentous environment (Fig. 2G, *t* = 35 s). These filament bends persist over time due to kinetic trapping by surrounding filaments (dashed circles in Fig. 2G, *t* = 70 s), given that the filament polymerization rate far exceeds the timescale of bending energy relaxation. The high-curvature filament shapes serve as templates for the assembly of additional filaments in the vicinity (Fig. 2G, *t* = 105 s), leading to spatially correlated bending, as evident from the spatial distribution of *θ*(***r***, *t*) in Fig. 2G.

### Nematic ordering and defect organization

To characterize the structure of the growing filament assembly, we analyzed the dynamics of nematic order in space and time. Local nematic order is associated with filament bending and the presence of topological defects within curved filamentous structures. The local orientation of filaments, represented by the orientation of each link, enables the construction of a nematic field for the network. It is evident from our simulations that regions with pronounced bending contained positive half-integer defects, while negative half-integer defects were more prevalent in areas of lower local density (Fig. 3A).

**Fig. 3.**
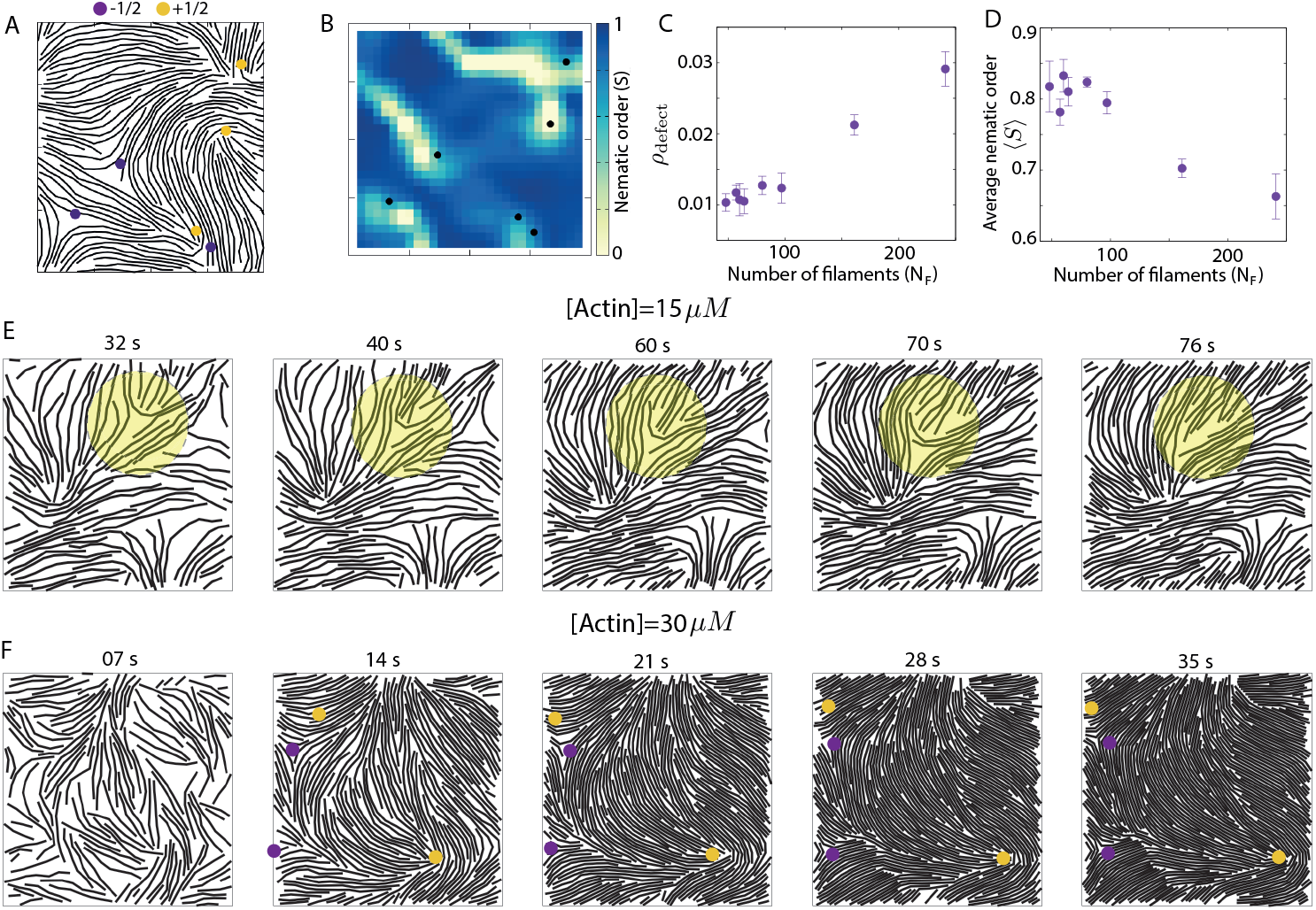
Nematic alignment and topological defects in filament assembly. (A) A representative configuration of a filament assembly, showing the distribution of half-integer defects. +1/2 defects are shown by solid yellow circles and -1/2 defects are shown in purple. (B) Spatial profile of local nematic order *S* for the filamentous network shown in panel (A). The black dots represent the position of half-integer defects. The yellow regions of ∼ 0 nematic order around the defects indicate the defect core size. (C) Defect density as a function of the number of filaments. (D) Spatially averaged nematic order *S*, as a function of the number of filaments. (E) Time-lapse sequences showing the evolution of actin filament configurations during growth at an actin concentration of 15 *μ*M. The shaded yellow region highlights an example of structural relaxation, where filament bending gradually relaxes over time, resulting in the disappearance of the defect. (F) Time-lapse sequences illustrating the configurations of actin filaments at a high actin concentration of 30 *μ*M. In this scenario, topological defects are long-lived as filament bends are unable to relax due to high crowding. In simulations for panels A-E, we chose [actin]=15 *μM*. For panels A-B, *N*_*F*_ = 64(*L* = 15 *μm*). For C and D, *N*_*F*_ = 97(*L* = 10 *μm*) and the results are averaged over 8 independent simulations.

Throughout the network, the depth and core size of these defects exhibited variations and were particularly localized in regions with correlated bending (Fig. 3B). These defects, indicative of higher energy bent structures, arise from the forces exerted by the growing filament barbed ends against neighboring filaments. The stability of the defects over time suggests a kinetic trapping phenomenon, wherein the filament bends are unable to relax due to crowding. Intriguingly, numerous instances of structural relaxations in regions of high filament curvatures were observed, leading to the subsequent annihilation of defect pairs (see Fig. 3E). These structural relaxations are mostly observed in long filaments at low and intermediate actin densities, when the timescale of growth is comparable to that of elastic relaxation. At high actin densities, structural relaxations are suppressed and filaments are permanently trapped in high curvature morphologies (see Fig. 3F and Movie S5).

We find that the density of half-integer defects (*ρ*_defect_), encompassing both +1/2 and -1/2 defects, increased with increasing filament number density (Fig. 3C), resulting in a corresponding decrease in the average nematic order (Fig. 3D). The decrease in filament length with increasing filament number contributes to the reduction in nematic order, where longer filament assemblies exhibit a globally high nematic order whereas shorter filaments possess many defects (see Fig. 2A, Fig. 3C-D).

### Increased filament density promotes correlated bending

The abundance of actin proteins plays a crucial role in regulating F-actin length and the resulting network morphology. To investigate the influence of actin density on network structure, we examined the growth and self-organization of F-actin networks across different values of total actin density. Elevated actin density, indicative of a larger G-actin pool, leads to the assembly of longer filaments at a given nucleator density, since 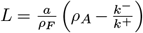. This, in turn, increases the F-actin density, leading to alterations in network morphology. As shown in Fig. 4A, higher actin density results in more extensive regions of correlated and heightened filament bending (also see Movie S5).

**Fig. 4.**
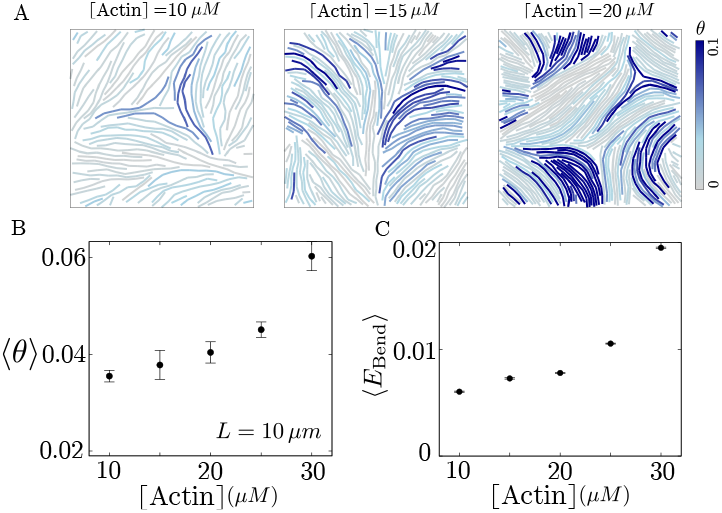
Effects of actin density on filament mechanics. (A) Filament configurations for different actin densities. (B) Average filament bending at the same filament length (10 *μm*) with increasing actin density. (C) Average bending energy at the same filament length (10 *μm*) with increasing actin density. Here *L* = 10 *μm* for all actin densities (i.e., different *N*_*F*_ values) and the results are averaged over 8 independent simulations.

Increasing filament density promotes interactions among filaments, leading to the formation of more kinetically trapped structures and consequently amplifying overall bending within the network (Fig.4B). Notably, there is an observed increase in the average bending energy per filament, *E*_Bend_, with the increase in actin density (Fig.4C). This underscores the discernible impact of crowding on network morphology, emphasizing the role of actin density in regulating filament curvature and alignment.

### Filament crosslinking amplifies kinetic trapping

Actin crosslinking proteins play a crucial role in maintaining the architecture of actin networks within cells. To understand impact of filament crosslinking on the evolving structure of a network, we simulated actin network growth under varying crosslink densities. Crosslinkers are expected to impose additional constraints and interactions among filaments. Indeed, as the crosslink density ([Xlink]) in the network is increased, we observed increased kinetic trapping of high filament curvatures, leading to an increase in average filament strains (Fig. 5A-C & Movie S6).

**Fig. 5.**
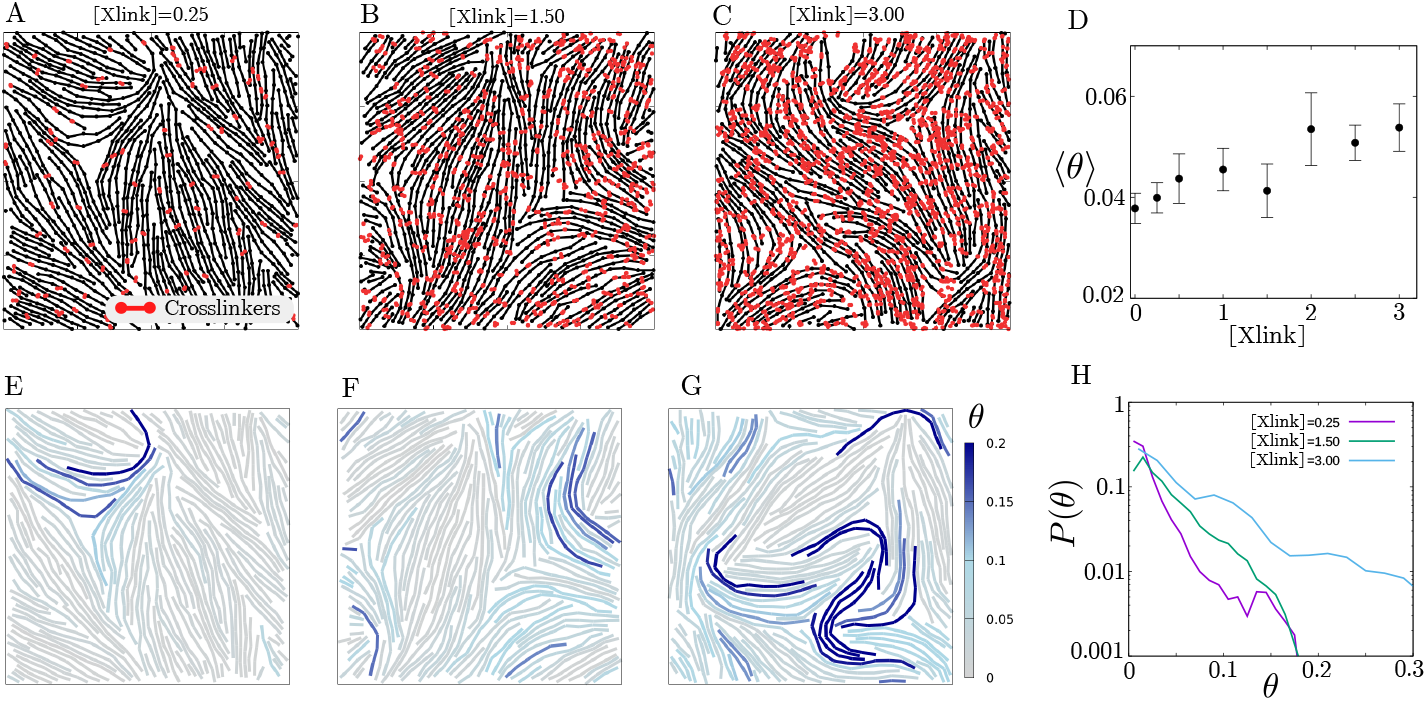
Crosslinking enhances filament bending. (A-C) Filament configurations for different crosslinker densities. The crosslinker is plotted in red. (D) Average filament bending with increasing crosslinker density. (E-G) Filament configurations coloured by filament bending for different crosslinker densities (same as A-C panels). (H) Probability distribution *P* (*θ*) of filament bending (*θ*) for different crosslinker densities. Here [actin]=15 *μM* and *N*_*F*_ = 97(*L* = 10 *μm*) and the results are averaged over 8 independent simulations.

To quantitatively assess the impact of crosslinking on collective filament bending, we computed the average compressive strain, ⟨*θ*⟩, which consistently increased with increasing crosslinker density (Fig. 5D). In addition to the elevated compressive strain, crosslinking facilitated the formation of larger regions of spatially-correlated filament bending (Fig. 5E-G). These alterations in network morphology became more apparent in the spatial distribution of strain *θ*, represented by the probability density *P* (*θ*). With an increasing crosslinker density, we observed a widening of the *P* (*θ*) distribution, indicating heterogeneity in filament bending throughout the network (see Movie S7), with the presence of some highly bent structures. Thus, variations in crosslinker density have the potential to control the emerging morphology of the developing actin network, leading to increased compressive strains and topological defects within the network.

## Discussion

In this article, we studied how the interplay between filament growth, bending elasticity and crowding regulates the emergent morphology of growing cytoskeletal networks. In particular, we found that filament polymerization in crowded environment induces correlated bending, forming high-curvature filamentous structures. These high curvature filaments lead to defects in nematic alignment that are kinetically trapped for extended period. Structural relaxation is observed for moderate actin densities and for long filaments where the timescale of bending relaxation competes with growth. While these findings are presented for filaments with physical properties resembling actin filaments, the implications extend to other cytoskeletal filament assemblies, including microtubules, intermediate filaments and FtsZ.

In cellular contexts, actin filament nucleation is typically regulated by nucleation-promoting factors and nucleators like formin and arp2/3, with spontaneous nucleation playing a negligible role (30, 31). The introduction of spontaneous nucleation may introduce significant heterogeneity in filament length distribution (29), a factor that is not considered in our study. We demonstrate that the precise regulation of filament length, occurring over the timescale of hours, is achievable when the filaments are nucleated within a sufficiently large actin monomer pool.

The limiting monomer pool enables us to regulate filament density and length by adjusting filament nucleation, replicating behaviors observed *in vitro* experiments (22). Specifically, we demonstrate that the network morphology undergoes significant changes depending on filament length, which is controlled solely by the number of nucleators seeded in the simulations. In networks with long filaments and moderate actin densities, we identify regions characterized by correlated and highly bent filaments, leading to the nucleation of long-lived topological defects. Tuning filament length and density allows for the tuning of nematic order and defect organization within the network. These morphological properties carry broader implications, potentially influencing how molecular motors generate forces within such dynamic networks. It’s important to note that the collective bending patterns and the emergence of high energy structures like topological defects are not driven by active motility, as observed in actin motility assays (32–34). Instead, the dynamics solely stem from filament growth, where filament barbed ends exert force on neighboring filaments during their growth, leading to the self-organization of filaments into a network.

Beyond nucleator density, both total actin density and the presence of crosslinker proteins play important roles in shaping the emergent structure of the growing network. An increase in actin density and filament crosslinking induce additional interactions among filaments, resulting in pronounced filament bending, extended regions of correlated bending, and a higher occurence of topological defects. These morphological changes can significantly impact various aspects of force generation and the mechanical properties of cytoskeletal networks. Notably, topological defects in cytoskeletal networks have been implicated in the regulation of key biological processes, including morphogenesis (35, 36). Exploring how molecular motors influence the organization of cytoskeletal networks with topological defects is an exciting avenue of future research.

## Methods

### Stochastic model for single filament growth

We use the Gillespie algorithm (37) to simulate the stochastic growth of multiple filaments from a shared pool of monomers. At any time *t* the algorithm uses two random variables drawn from a uniform distribution (*r*_1_, *r*_2_ ∈*𝒰* (0, 1)), and the instantaneous propensities for all of the possible reactions to update the system in time according to the defined growth law. The propensities of the relevant reactions, i.e., the assembly and disassembly rates of the *i*^*th*^ actin filament (consisting of *n*_*i*_ monomers) are given by 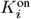 and 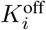 respectively. For the limiting pool model these propensities are functions of monomer pool size (*N*) and structure size (*n*_*i*_),

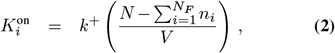

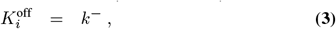

where we are considering the growth of *N*_*F*_ filaments from a shared monomer pool and *k*^+^ and *k*^−^ are the rate constants for assembly and disassembly from the ends of the actin filament. We use the simplification of considering combined rates of assembly 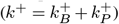 and disassembly 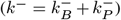 rather than modeling barbed ends and pointed ends separately (Fig. 1A). We also do not consider the hydrolysis of the filament-bound monomers for simplicity.

The Gillespie algorithm computes the time for the next reaction at *t* + *τ* given the current state of the system (i.e., the propensities for all reactions) at time *t* where *τ* is given by-

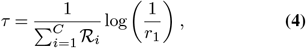

where ℛ_*i*_ is the propensity of *i*^*th*^ reaction and *C* is the total number of all possible reactions. The second random variable *r*_2_ is used to select the particular reaction (*j*^*th*^ reaction) that will occur at *t* + *τ* time such that

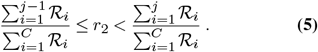

The condition for the first reaction (*j* = 1) is 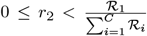. The two steps defined by Eq. 4 and Eq. 5 are used recursively to compute the growth dynamics over time.

### Effective model of single filament growth

An equivalent deterministic description of the limiting pool model described above will look like:

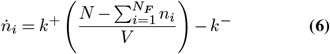

for the *i*^*th*^ filament where *i* ∈ (1, .., *N*_*F*_). Length of the *i*^*th*^ filament is determined as *L*_*i*_ = *an*_*i*_, where *a* is the size of a monomer. This set of ODEs result in a set of under-determined equations when the steady-state filament length (given by 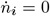) is sought for. The possible steady-state solutions are given by a constraint 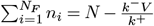. In the case of a large monomer pool, the filaments have small length fluctuations and they acquire similar length (*n*), i.e., 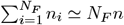. This approximation based on the results of stochastic growth (Fig. 1B&C) leads to an approximate equation of length dynamics as shown in Eq. 1.

### Agent-based simulation of growing filaments

The agent-based simulations were conducted using the AFINES package (18) to explore the morphology of growing networks involving F-actin and cross-linkers. We systematically varied nucleator density, actin density and cross-linker density, to understand the effects of growth and network composition on the emergent dynamics of network morphology. In our model, actin filaments were modeled as polar worm-like chains, consisting of *N* + 1 beads connected by *N* harmonic springs and *N* − 1 angular harmonic springs. We incorporate volume exclusion interactions between the links of any two filaments within a cutoff distance *r*_ve_. The volume exclusion energy between two links is taken to be

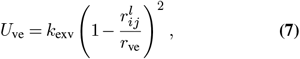

where the parameter *k*_exv_ controls the strength of volume exclusion interaction and 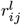 is the distance of closest approach between the *i*^th^ and *j*^th^ links. The potential energy of an actin filament is given by

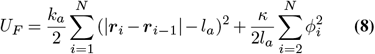

where ***r***_*i*_ represents the position of the *i*^*th*^ bead on a filament, *ϕ*_*i*_ is the angle between the *i*^*th*^ and *i* − 1^*th*^ links, *k*_*a*_ is the stretching force constant, *κ* is the bending modulus, and *l*_*a*_ is the equilibrium length of a link. The persistence length (*L*_*p*_) of the filament was derived as *κ/k*_*B*_*T*, where *k*_*B*_ is Boltzmann’s constant and *T* is the temperature (see Eq. 10 below). The filament starts with a single link and grows from the barbed end until the barbed end link becomes 2*l*_*a*_ when a new bead is introduced at the middle of the link. Link growth is implemented by growing the rest length of the harmonic spring (corresponding to the barbed end link). Growth stops when the filaments reach a steady-state length determined by the approximate theory (Eq. 1).

Cross-linkers were modeled as Hookean springs, each with two ends (heads 1 and 2) capable of binding to and unbinding from filaments. The potential energy of a single cross-linker was expressed in

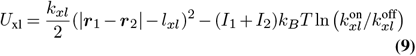

where ***r***_1(2)_ is the position of head 1(2), *I*_1(2)_ is 1 if head 1(2) is bound and 0 otherwise, and 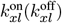 is the rate constant for binding (unbinding). When a cross-linker is bound, it moves with the filament to which it is attached. The tensile force *F*_*xl*_ stored in the cross-linker with a bound head at position *r*_*xl*_ is propagated to neighboring filament beads using the lever rule (18). Binding and unbinding events follow a Monte Carlo procedure designed to satisfy detailed balance. The time evolution of the network was simulated using Brownian dynamics, with the position of an actin bead or cross-linker head at each timestep generated from the previous one using

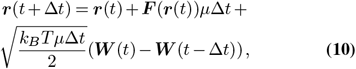

where ***F*** (***r***(*t*)) is the gradient of the potential of the particle (Eqs. 8 and 9), ***W*** (*t*) is a vector of random numbers drawn from the standard normal distribution, and we used the Stokes relation *μ* = 1*/*(6*πνR*) in the damping term, where *R* is the size of the particle, and *ν* is the dynamic viscosity of its environment. We simulated a 2D system of size 20 *μ*m×20 *μ*m at constant temperature *T* ∼ 300K. A list of important model parameters and their values for the simulations used in the paper is provided in Table 1. To limit boundary effects associated with the finite size of the simulation, periodic boundary conditions were used. For more details and a complete list of parameters please refer to Freedman *et al* (18).

**Table 1.**
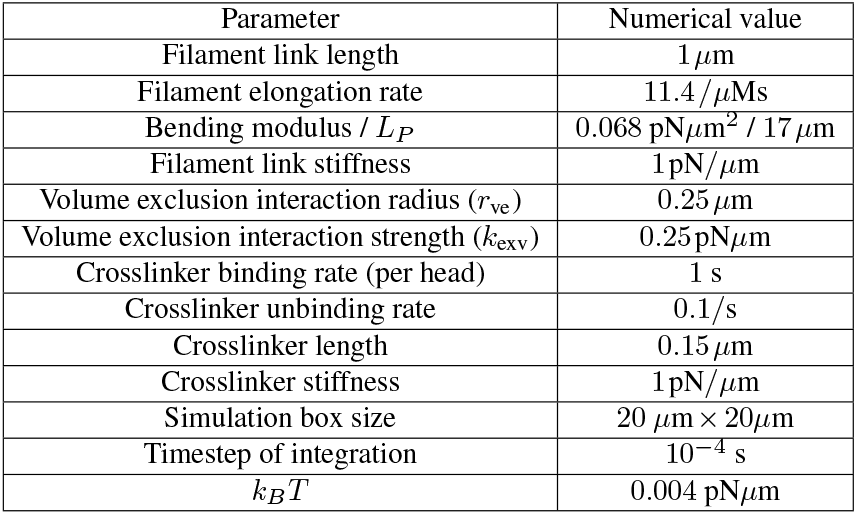
Default parameter values used in agent-based simulation with AFINES.

### Analysis of filament and network morphology

To quantitatively assess the morphological changes in the resulting network due to filament growth, we evaluate the extent of filament bending by calculating the compressive strain on each filament as

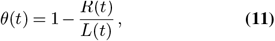

where *R*(*t*) is the end-to-end distance of a filament and *L*(*t*) is the filament contour length at time *t* (Fig. 2B). We present the temporal evolution of bending by calculating the average compressiver strain per filament (⟨*θ*(*t*) ⟩). We evaluate the extent of bending with changing parameters by calculating a mean compressive strain (⟨*θ⟩*) averaged over time (in steady-state, i.e., last few time points) and over the ensemble.

We also calculate the nematic order in the network and the number of topological defects. The nematic director field ***n***(***r***, *t*) was created by interpolating direction vectors (using weights exponentially decaying with distance) from the local orientation (*?*) of filament segments in a small box. Then the local director field was used to calculate ***n***(***r***, *t*) = (*cos*(*?*), *sin*(*?*)). The topological defects were detected by using the winding angle approach (38).

## Supplementary Movies

### Movies S1-S4

These movies show the time evolution of growing filament networks with actin density 15 *μ*M and different nucleator numbers such that the steady state filament lengths are 20 *μ*m (Movie S1), 15 *μ*m (Movie S2), 10 *μ*m (Movie S3) and 4 *μ*m (Movie S4), respectively. The filaments are colored by their respective compressive strain *θ* values, as marked in the colorbar. The parameters of these simulations correspond to the four panels in Fig 2A.

### Movie S5

This movie shows the assembly of a growing filament network at high F-actin density, resulting in a crowded environment characterized by highly bent filaments that are kinetically trapped and remain highly stable without undergoing structural relaxation. The actin density here is 30 *μ*M and the steady state length is 17 *μ*m. This movie corresponds to the time-lapse snapshots presented in Fig 3F.

### Movie S6

This movies shows the time evolution of a growing network with crosslinkers (shown in red) at density [Xlink]=3, actin density 15 *μ*M and steady state filament length 10 *μ*m. The crosslinkers are shown in red. This movie corresponds to the network configurations presented in Fig. 5C.

### Movie S7

This movies shows the time evolution of a growing network with crosslinkers (not shown) at density [Xlink]=3, actin density 15 *μ*M and steady-state filament length 10 *μ*m. The filaments are colored by their compressive strain *θ* values as marked in the colorbar. This movie corresponds to the network configurations presented in Fig. 5G.

## Acknowledgements

SB acknowledges support from the National Science Foundation (NSF MCB-2203601) and the National Institutes of Health (NIH R35 GM143042). MM acknowledges ARO MURI W911NF-14-1-0403, the National Institutes of Health (NIH) R01 1R01GM126256, and National Institutes of Health (NIH) U54 CA209992.

